# Genome-Wide Markers Predict Metribuzin Tolerance in Southern Soft Red Winter Wheat

**DOI:** 10.64898/2026.06.28.733875

**Authors:** Julio Sellani, Hugo Anzueto, Kelly Arcenaux, Paul “Trey” Price, Gina Brown-Guedira, Stephen Harrison, Noah DeWitt

## Abstract

Metribuzin is a versatile herbicide effective against various annual grasses and broadleaf weeds found in wheat fields. However, it can cause foliar damage to wheat, impacting plant health and yield. A clearer understanding of the genetic architecture associated with metribuzin tolerance is necessary to guide marker-based breeding strategies. This study evaluated 351 historic Gulf Atlantic Wheat Nursery (GAWN) wheat breeding lines representative of southern US soft red winter wheat (SRWW) germplasm. Field trials were conducted at Winnsboro (WN) and Baton Rouge (BR), Louisiana, in 2016 and 2017. Metribuzin was applied at specific growth stages, and tolerance was assessed based on visual foliar damage. Genomic data from 6,252 filtered single nucleotide polymorphism (SNP) markers were used to estimate narrow-sense heritability, conduct genome-wide association (GWAS), and assess genomic prediction accuracy using genomic best linear unbiased prediction (GBLUP). Broad-sense heritability ranged from 0.54 to 0.69 within environments and reached 0.77 across environments, while narrow-sense heritability ranged from 0.35 to 0.47, indicating moderate additive genetic control. No SNP surpassed the significance threshold, but genomic prediction (GP) showed moderate to strong predictive ability (PA) across environments, with the highest accuracy (r = 0.62) observed between BR17 and WN17. These results indicate that metribuzin tolerance in SRWW is primarily controlled by multiple small-effect loci and that GS provides a more effective breeding strategy than marker-assisted selection for improving tolerance in southern wheat germplasm.

## 1 INTRODUCTION

Wheat (Triticum aestivum L.) is one of the world’s major food crops and provides nearly 20% of calories consumed, making it the third most important crop globally after corn and rice (FAOStat https://www.fao.org/faostat/en). Wheat production is negatively affected by biotic and abiotic factors that include weather, diseases, insect pests and weeds. The competition between weeds and crops results in a reduction of grain yield, attributed to the loss of space, nutrients, water, and light for plant growth and reproduction (Siddiqui et al., 2010). High weed pressure decreases wheat tillering and vigor, leading to a significant reduction in grain yield. Notably, heavy weed infestation severely limits wheat productivity, leading to a more than 48% reduction in potential wheat yield (Fahad et al., 2015). Therefore, the easiest way to prevent yield loss is to use herbicides to control weeds while the crop grows. Some crops are tolerant of specific herbicides and survive the application with little or no damage (Bhullar et al., 2017). Choosing the appropriate herbicide is important to avoid damage. The development of wheat varieties tolerant to specific herbicides can broaden growers’ weed management options and reduce crop injury associated with herbicide applications (Runyan et al., 1982; Baker and Peeper, 1990).

Metribuzin (4-Amino-6-(1,1-dimethylethyl)-3-methylthio-1,2,4-triazin-5(4H)-one) is versatile and highly effective for weed management and provides control of numerous important annual grasses and broad leaf weeds in wheat. Metribuzin is a common triazine herbicide that is highly soluble (1200 mg/L) and is absorbed into the roots by diffusion when it is applied to the soil, being translocated via xylem to the shoots. Metribuzin penetrates the symplast but is not retained and leaches back into the apoplast (WSSA, 2014). Applied post-emergence, metribuzin is absorbed through foliage and roots and exhibits limited translocation within the plant (Shaner, 2014). Metribuzin inhibits electron transport by binding to the D1 protein of the PSII reaction center, blocking electron transfer to plastoquinone. Inhibition of PSII electron transport prevents the conversion of light energy into chemical energy and ultimately disrupts photosynthesis (Trebst, 2007). By blocking electron transport, the electrons react with chlorophyll, creating triplet chlorophyll which reacts with the ground stage oxygen creating singlet oxygen; these two components lead to a chain reaction of lipid peroxidation that degrades the chlorophyll, carotenoid, and cell membrane, making the cell lose its integrity and start to dehydrate (WSSA, 2014). Metribuzin has been adopted in some production systems as an alternative herbicide option where concerns regarding atrazine use have increased. (Graymore et al., 2001).

Metribuzin is commonly used on wheat fields in the Southeast United States because it is effective and inexpensive. In addition to its use in Southeast, metribuzin remains an important weed management tool in major wheat-producing states such as Kansas, Oklahoma, Texas, and Kentucky, and is also widely spread in soybean, barley, potato, and sugarcane production throughout the United States (Albaugh, 2020; Singh et al.2025). Baker and Peeper (1990) found that wheat cultivars show differences in their response to metribuzin applications, depending on the dose and environment. In prior field screening at two locations in Louisiana, wheat lines showed substantial damage at recommended rates that may cause yield loss. The development of elite metribuzin resistant lines would facilitate wheat production in the South by reducing input costs, but phenotyping for resistance is complicated and resource intensive. Studies have demonstrated metribuzin tolerance in wheat lines (Runyan, 1982). Pilcher et al. (2017) concluded that the available information regarding the genetics of metribuzin tolerance inheritance in wheat remains limited. They reported moderate levels of heritability in both the broad-sense (H = 0.52) and narrow-sense (h^2^ = 0.23) for tolerance to metribuzin, and tolerance was influenced by the combined action of multiple alleles. Garcia-Baudin et al. (1990) reported that durum cultivars (Triticum turgidum var. durum L) ‘Anton’ and ‘Nita’, exhibit tolerance and susceptibility to metribuzin, respectively. That wheat demonstrates different levels of metribuzin tolerance, and that this variation is heritable, suggests that marker-based breeding strategies may be utilized to identify tolerant varieties with less field-based phenotyping.

Marker-based predictive breeding approaches may enable selection for tolerance but require a more detailed understanding of the genetic architecture for this trait in wheat. Strategies for marker-based breeding depend on the genetic architecture of the trait of interest, and establishing an understanding of the feasibility and optimal approach is the focus of this study. Previous work has investigated the genetic basis of metribuzin tolerance in other crop species. In general, it has been found to be a heritable, quantitative trait, with multiple genomic regions contributing to phenotypic variation in wheat (Kurya et al., 2022). Identification of markers closely linked to major metribuzin tolerance quantitative trait loci (QTL) would improve selections for this trait in wheat breeding, allowing for marker-assisted selection (MAS) that could speed and improve the efficiency of breeding for metribuzin tolerance. The introduction of genomic selection presents an alternative route whereby genome-wide markers can be used to predict quantitative traits to make selections in breeding programs, even absent the presence of major QTL. To determine the best approach for utilizing markers in breeding for this trait, the present study focuses on the identification of the genetic architecture of metribuzin tolerance and efficacy of genomic prediction methods in soft red winter wheat germplasm representative of current Southeast US breeding lines.

## 2 MATERIALS AND METHODS

### Genotypic Data and Quality Control

Three hundred and fifty-one diverse wheat genotypes were used in the study, consisting of varieties and breeding lines drawn from the Gulf Atlantic Wheat Nursery (GAWN) from 2008-2014. The GAWN is a late yield stage trial of advanced breeding lines from the SunGrains® collaborative breeding program, which consists of the small grains breeding programs at the University of Arkansas, University of Florida, University of Georgia, Louisiana State University, North Carolina State University, Clemson University, and Texas A&M University, and external partner Virginia Tech.

The 351 genotypes were selected based on seed availability and their adaptation to the short vernalization environment of Louisiana. The USDA Eastern Regional Small Grains Genotyping Lab generated SNP data using the 90K SNP chip, containing 81,578 SNP markers. Genotype data were read from a VCF file and converted into numeric genotypes genotype codes appropriate for diploid individuals. For each SNP, lines homozygous for major alleles were coded as 0, heterozygous as 1, and homozygous for the minor allele as 2. The filtering of the data deleted markers with missing data of greater than 10% and minor allele frequency less than 5%. Filtering was followed by thinning of the marker matrix based on a linkage disequilibrium threshold of 0.80. The final data set had 6,252 markers.

### Field Evaluations

Trials were planted at the Louisiana State University Agricultural Center (LSU AgCenter) Central Station Ben Hur Research Farm in Baton Rouge (BR), LA and at the Macon Ridge Research Station in Winnsboro (WN), LA, in November 2015 and November 2016. A randomized complete block design with 2 replications and 351 plots per replication was used for each environment. Each plot consisted of two rows, each 102 cm long, with 38 cm spacing between the rows. Pre-plant phosphorus, potassium, and sulfur were applied as necessary at each location based on a soil sample analysis at the LSU AgCenter Soil and Plant Testing Laboratory. No pre-plant nitrogen was applied. Plots were top dressed with 79 kg N/ha in the spring with 28-0-0-2 or 32-0-0 liquid fertilizer.

The outer two rows on each side of every 54 rows wide range were planted with filler wheat to act as borders to prevent any effect from the lack of competition of exterior rows. As a pre-plant weed control measure, the fields were sprayed with glyphosate in the fall. Alleys were sprayed as needed in spring with glyphosate using a covered sprayer behind a four-wheel all-terrain vehicle to clear space between the ranges of the rows. Metribuzin was applied at a rate of 527 grams per hectare (10 oz/acre of 75DF) of active ingredient when the wheat was at Feeke’s growth stage 4-5 on 3/04/2016 and 3/03/2017 in WN and 2/17/2017 in BR. The application was made with a four-wheel all-terrain vehicle driven down the open alleys at 8km/hr with a spray boom of 5.33 m long that covered 5.55 m width (2 ranges of plots). An electric pump was used to pressurize the herbicide solution and a standard liquid regulator with a pressure gauge was used to regulate the flow. Metribuzin was applied using TeeJet® (Teejet Technologies, LCC, Illinois) 11003 spray nozzles and a herbicide volume of 140 liters per hectare.

The field was inspected every other day after herbicide application to note the date when the most sensitive genotype showed damage. The ratings were taken 14 days after application using a visual scale from 0-9 (0 indicates no damage, 1-3 resistant, 4-6 moderately resistant/sensitive, 7-8 sensitive, and 9 indicates complete plant death). The data was collected in BR (2017) and in WN (2016, 2017) as satisfactory crop stand was not established in Baton Rouge in 2016 due to heavy rain.

### Estimation of Genotypic Values for Metribuzin Tolerance

R version 4.2.2 (2022-10-31) was used to obtain Best Linear Unbiased Estimates (BLUEs) for genotypes, estimate variance components assessed using the R package “sommer” (Covarrubias-Pazaran, 2016), and to perform genomic prediction the R package “rrBLUP” (Endelman, 2011).BLUEs for genotype foliar damage were estimated using linear models that fit separately for each environment (within-location) and for the combined dataset (across-location). For all models, BLUEs were calculated using the following linear model:

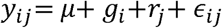

where *y_ij_* is the observed foliar damage for genotype *i* in replicate *j*, *µ* is the overall mean, *g_i_* is the fixed effect of genotype, *r_j_* is the fixed effect of replication, and 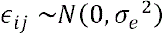 is the residual error. This model was applied consistently to each site-year (WN16, WN17, BR17).

For the across-location dataset (ALL), a modified model was run fitting site-year as fixed effects, and fitting replication nested within site-years:

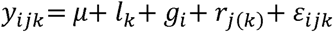

where *y_ijk_* is the observed foliar damage score for genotype in replicate *j* within site-year *k*, *µ* is the overall mean across all site-years and genotypes, *l_k_* is the fixed effect of site-year (environment), capturing environmental variation across years and sites, *g_i_* is the fixed effect of genotype, representing genetic contributions to foliar damage response, *r_j(k)_* is the fixed effect of replication nested within site-year, accounting for block structure within each environment, and 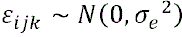 is the residual error term assumed to be normally distributed and independent. No genotype-by-environment interaction term was included in the combined model. Models were fit using the “mmer()” function from the “sommer” package, with predictions extracted based on the fitted fixed genotype effects.

To visualize and statistically compare BLUEs across breeding programs, Tukey’s Honest Significant Difference (HSD) test was used to identify significant differences in mean metribuzin tolerance among breeding programs within each environment. The test was applied following an analysis of variance (ANOVA), where the breeding program was treated as the fixed factor and BLUE values as the response variable. Tukey’s HSD controls the family-wise error rate while making all pairwise comparisons and assumes that residuals are normally distributed with equal variances. The compact letter display (CLD), which groups breeding programs with similar performance, was generated using the “multcompView” package in R. This package takes the results of Tukey HSD and assigns letter groupings that summarize pairwise significance comparisons. This allowed for an intuitive visualization of statistical differences in BLUEs across breeding programs within each environment.

### Heritability Estimation

Heritability estimates were calculated in R using traditional broad-sense, Cullis-adjusted broad-sense, and narrow-sense approaches across individual site-years (WN16, WN17, BR17) and for the combined dataset (ALL). Traditional broad-sense heritability (H) was estimated using the same models as detailed above, except treating the genotype as a random effect instead of fixed. For the within-site-year models a simple random intercept model was used. Broad-sense, entry-mean heritability was calculated using the equation:

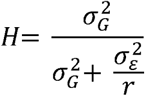

where 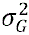 is the genotypic variance, 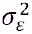 is the residual variance, and r is the number of replications.

Cullis heritability H_Cullis_ was estimated following the approach of Cullis et al. (2006), using the “lmer()” function from the “lme4” package. Within each site–year, a linear mixed model with genotype as a random effect was fitted to generate Best Linear Unbiased Predictions (BLUPs) for the genotypes. For the combined dataset, the model included site–year as a fixed effect and replication nested within site–year as a random effect:

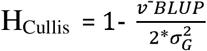

where 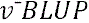 is the average prediction error variance (PEV) of the pairwise differences between genotypic BLUPs and 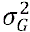 is the genotypic variance extracted from the model.

Narrow-sense heritability *h*^2^ was estimated using a genomic best linear unbiased prediction model (GBLUP), implemented via the “mixed.solve()” function from the rrBLUP package. The model incorporated a genomic relationship matrix K built from SNP marker data using the Van Raden (2008) method as implemented in the rrBLUP (Endelman, 2011), function “A.mat()”. Additive genetic variance 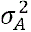 and residual variance 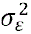 were extracted from the GBLUP model, and for each environment marker-based heritability was calculated as:

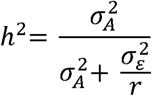

where *r* is the number of replications for that environment. Because this estimate is derived from a genomic relationship matrix and assumes all marker effects are additive, it represents a marker-based upper bound on the true narrow-sense heritability, reflecting the proportion of phenotypic variance captured by genome-wide SNPs under the additive model.

### Genome-Wide Association Study

A genome-wide association study (GWAS) was conducted to identify genomic regions associated with metribuzin tolerance using a standard SNP-based GWAS approach was used with the linear mixed model (Yu et al. 2006). Genome-wide association analysis (GWAS) was conducted using the GWAS() function from the rrBLUP package. The GWAS model used, where each SNP was tested individually using a linear model, has the form:

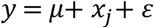

where *y* represents the BLUEs for foliar damage, *µ* is the overall mean, *x_j_* is the SNP genotype at locus *J* coded as dosage, and *ε* is the residual error. To account for genetic relatedness among individuals, SNP effects were tested within a linear mixed model of the form:

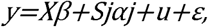

where yis the vector of genotype BLUEs for foliar damage, *X,β* includes an intercept where no principal components were fitted, *s_j_* is the vector of SNP genotypes for marker *j* coded as dosage, and *α_j_* is the fixed effect of SNP *j*. The random genetic effects were assumed to follow 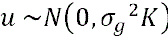, where K is the additive genomic relationship, kinship, matrix calculated using the “A.mat()” function from the “rrBLUP” package, and residuals assumed to follow 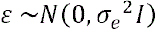. The analysis was performed separately for each set of BLUEs: within each environment (WN16, WN17, BR17) and the combined environment model (ALL), allowing assessment of genetic associations under different environmental conditions.

### Genomic Selection and Predictive Ability Assessment

Genomic prediction was performed using the Genomic Best Linear Unbiased Predictor (GBLUP) method to predict metribuzin tolerance across environments. The prediction model was implemented using the “mixed.solve” function from the “rrBLUP” package in R, incorporating relationship information using the same GBLUP model used for estimating narrow-sense heritability, except that y represents the vector of genotypic BLUEs instead of raw phenotypes. Correlations between observed means and predicted values were calculated to assess model fit. Predictive ability was assessed via five-fold cross-validation repeated over 40 iterations for each environment or combination of environments. For each fold, 20% of individuals were treated as unobserved, and GEBVs were predicted for those unobserved individuals using the model trained on the remaining data. Predictive ability was evaluated as the Pearson correlation between observed BLUEs and predicted GEBVs across folds.

### Phenotypic Classification and Environmental Consistency Analysis

To identify consistently resistant and susceptible lines, average foliar damage scores were calculated across the three tested environments (BR17, WN16, WN17), excluding missing values. Genotypes were ranked based on their across-location Best Linear Unbiased Estimates (BLUEs), with lower BLUE values indicating greater tolerance to metribuzin. The ten most resistant and ten most susceptible lines, as determined by these BLUEs, were selected for further visualization. These genotypes were categorized and displayed in a scatter plot, where each genotype’s BLUE value was plotted and color-coded according to its classification group (resistant or susceptible).

To evaluate the consistency of genotype performance across environments, pairwise Spearman rank correlation coefficients were calculated comparing BLUEs across locations to assess the strength and direction of relationships between lines rankings across site-years. The resulting correlation matrix was visualized as a heatmap, with cell values indicating the degree of agreement in relative tolerance rankings between each pair of environments.

## 3 RESULTS

### Heritability

Heritability estimates were calculated to assess the proportion of phenotypic variation attributable to genetic factors and thus repeatability of the metribuzin screening method across different environments. Broad-sense entry-mean heritability (H) was estimated using two different methods, a standard estimator using variance component ratios and the Cullis estimator. Narrow-sense heritability (h^2^) was also estimated based on additive genetic variance components (single and across-environment entry mean heritability). For broad-sense heritability, estimates were obtained both within individual site-years (single-environment entry mean heritability) and across all environments combined (across-environment entry mean heritability). The estimates are summarized in Figure 1. Broad-sense entry-mean heritability was highest when considering all environments combined (H = 0.77), which reflects genetic correlations among site-years and the increased precision in genotype value estimates due to a greater number of replications across environments. Among the individual site-years, the highest estimate was observed in WN17 (H = 0.69), and the lowest estimate was observed in WN16 (H = 0.54). Cullis heritability followed a similar pattern, with the overall estimate across environments being slightly lower (HCullis = 0.75). Among individual environments, WN17 exhibited the highest heritability across both estimators (HCullis = 0.69), while WN16 had the lowest (HCullis = 0.54).

**Figure 1.**
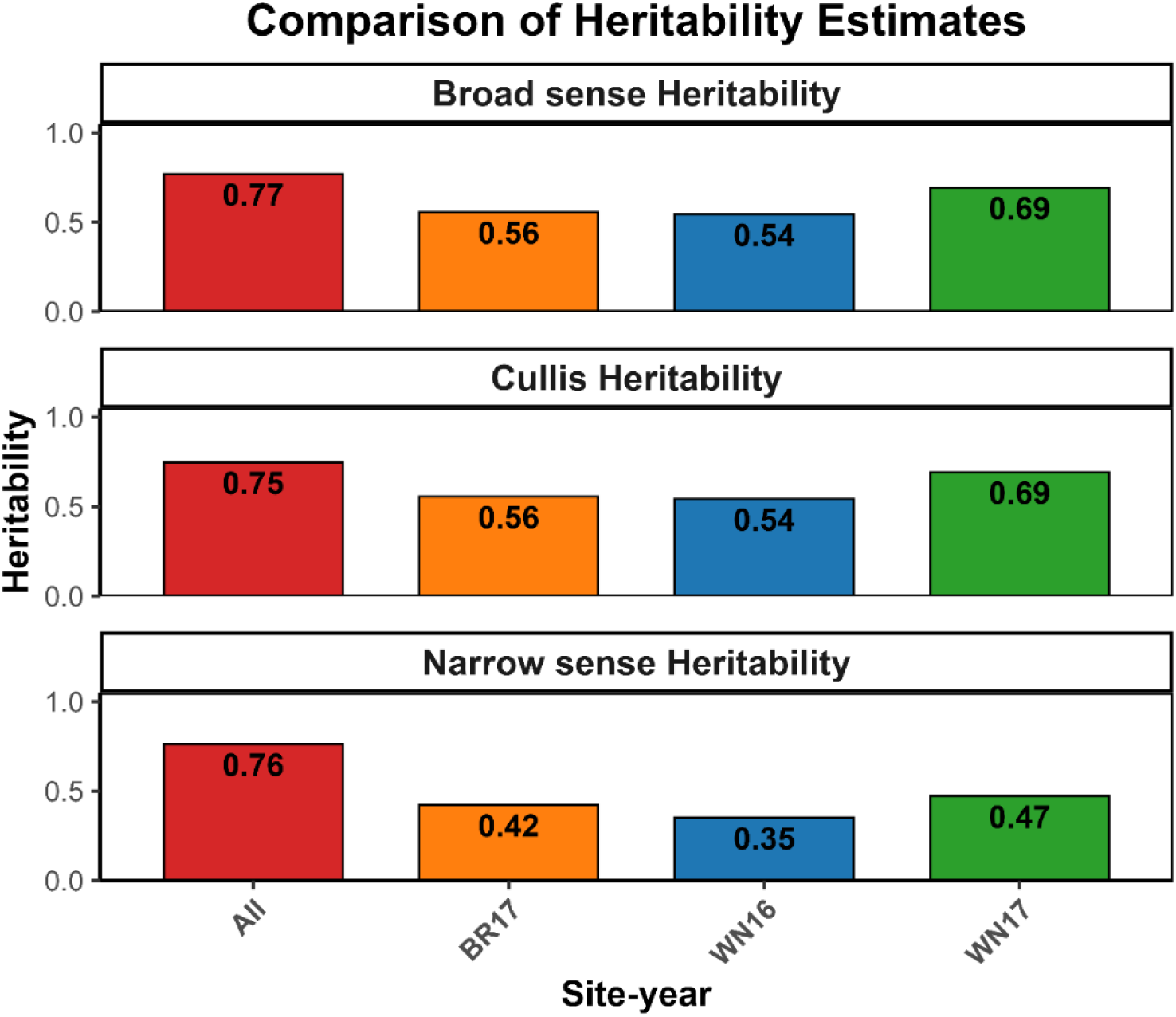
Broad-sense (H), Culli broad-sense (HCullis), and narrow-sense (h^2^) entry-mean heritability estimates for metribuzin tolerance across individual site-years (BR17, WN16, WN17) and the combined multi-environment dataset (All). Broad-sense heritability was estimated using variance components from mixed models treating genotype as random effect. Cullis heritability was calculated based on the average prediction error variance of genotypic BLUP differences. Narrow-sense heritability was estimated using a genomic relationship matrix in a GBLUP framework. The “All” category represents across-environment entry-mean estimates incorporating all site-years.

Estimates of narrow-sense heritability (h²) were generally lower than broad-sense and Cullis heritability estimates. Across all environments, entry-mean h² ranged from 0.35 to 0.76, suggesting that non-additive genetic effects, such as dominance and epistasis, contributed to trait expression. Among site-years, the highest narrow-sense heritability was observed in WN17 (h² = 0.47), while the lowest was in WN16 (h² = 0.35), indicating that the proportion of phenotypic variation explained by additive effects varied across environments. The combined analysis across environments (“All”) yielded the highest estimates for all three methods, emphasizing that integrating multi-environment data improves the precision of heritability estimation by reducing environmental noise. The high narrow-sense heritability in the combined dataset also indicates that additive effects are largely stable across environments, supporting the potential for effective genomic selection for metribuzin tolerance when multi-environment data are utilized.

### Genotypic Performance and Environmental Repeatability

To assess the consistency of metribuzin tolerance expression across environments, pairwise Spearman rank correlations were calculated using BLUEs from three site-years. The correlation (Supplementary Figure S1.) between environments BR17 and WN17 ρ = 0.61**, indicates that genotypic rankings for foliar damage were similar across these environments. In contrast, weaker correlations were observed between BR17 and WN16 (ρ = 0.41*), and between WN16 and WN17 (ρ = 0.43*).

Figure 2 visually captures both the average foliar damage scores and the environment-specific responses for the 10 most resistant and 10 most susceptible genotypes. The larger red and green dots represent average damage ratings across environments, while smaller dots represent genotype scores in each individual site-year (BR17, WN16, WN17). This multi-point visualization denotes those resistant lines (e.g., AR04006-1, AR05055-12-2) consistently exhibited low foliar damage across environments, reinforcing their stability and suitability as parents in breeding programs targeting metribuzin resistance. Eight of the ten most resistant genotypes originated from the Arkansas breeding program. In contrast, the most susceptible genotypes showed greater variability in metribuzin damage scores across environments. For example, genotype SCLA18084A1 had a damage rating of 3.3 in WN17 but 7.2 in WN16, indicating substantial environmental effects on susceptibility.

**Figure 2.**
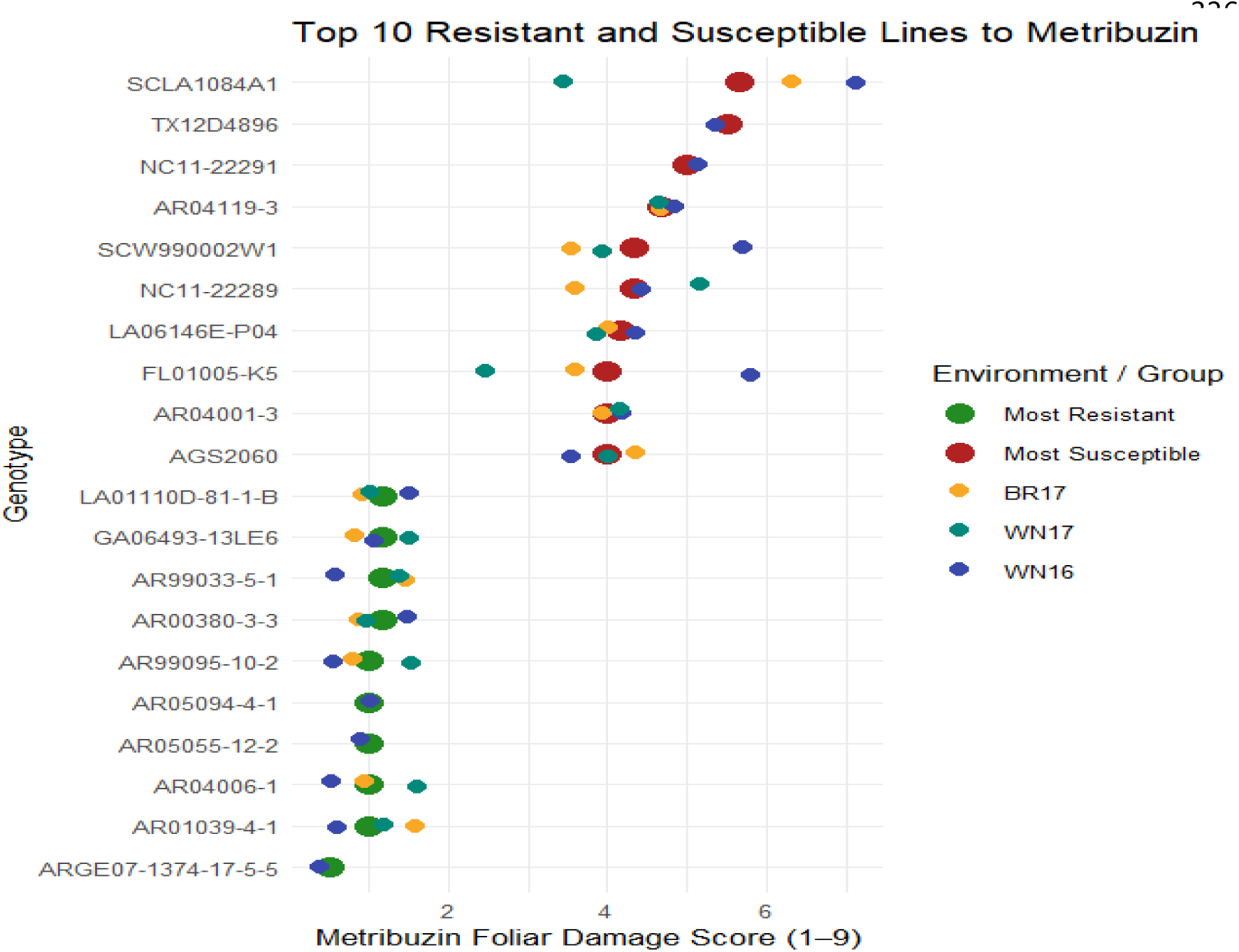
Average foliar damage scores across environments for the ten most tolerant (lowest scores) and ten most susceptible (highest scores) wheat genotypes. Large dots represent the across-environment mean, while smaller colored dots indicate individual scores within each environment (BR17 = gold, WN16 = blue, WN17 = teal). Genotypes are grouped and labeled by classification: “Most Resistant” (green) and “Most Susceptible” (red). Lower scores indicate greater tolerance (1 = no damage; 9 = complete chlorosis/necrosis).

### Regions Associated with Metribuzin Tolerance

To identify loci associated with metribuzin response, a genome-wide association study (GWAS) was conducted using the rrBLUP model (Figure 3). The combined analysis across all environments (ALL) revealed but no SNPs that exceeded a genome-wide significance threshold, consistent with a polygenic architecture for this trait.

**Figure 3.**
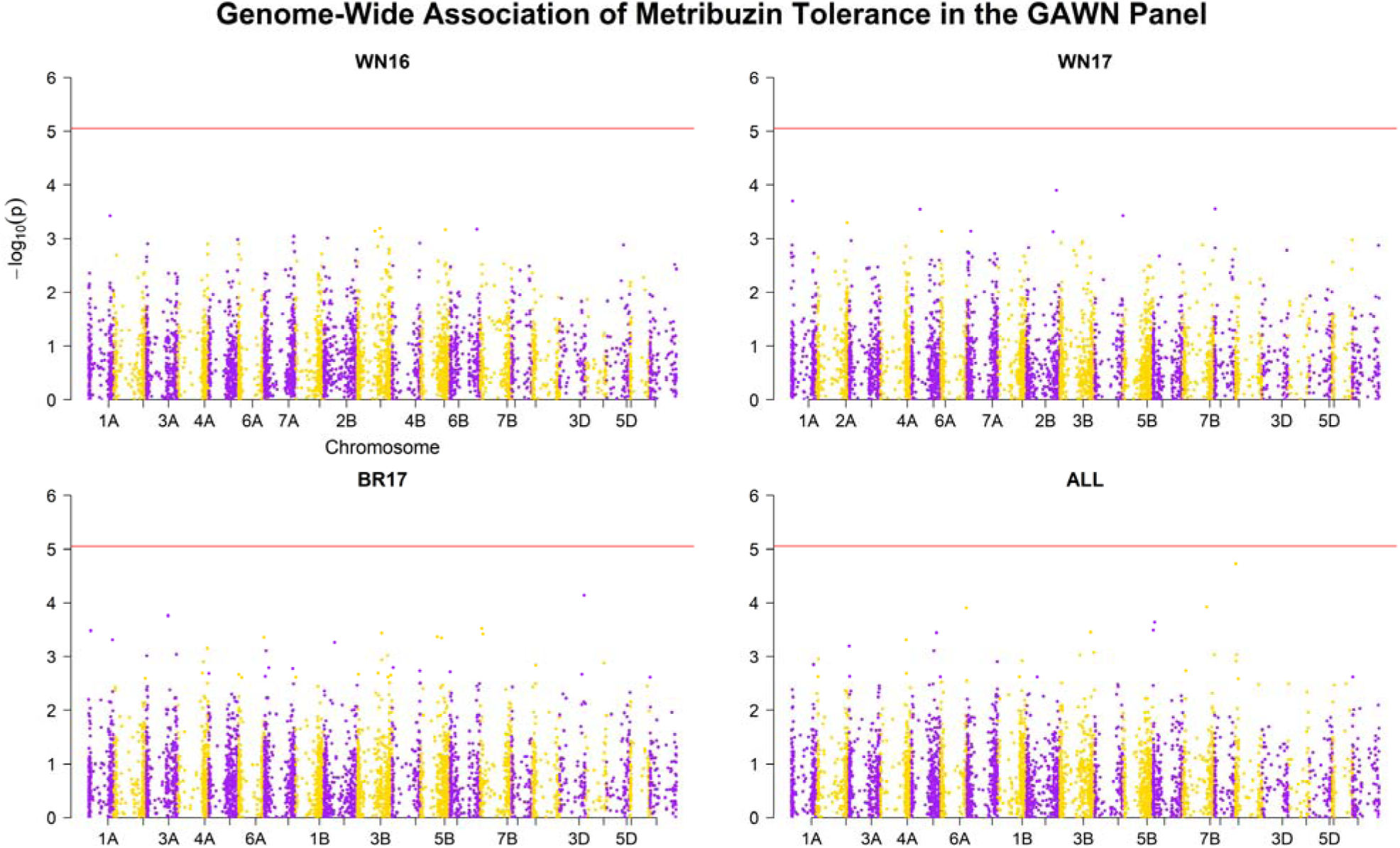
Manhattan plots illustrate genome-wide association results for metribuzin tolerance across three environments (WN16, WN17, BR17) and the combined dataset (ALL). Each dot represents a single SNP, plotted according to its physical position along the wheat genome (x-axis) and its statistical significance (-log□□(p-value), y-axis). Results for each environment are plotted separately. No SNP exceeded the genome-wide significance threshold based on a Bonferroni correction (α = 0.05; threshold = 5.05).

### Predictive Ability Within and Across Environments

To assess the effectiveness of genomic prediction for metribuzin tolerance, a predictive ability matrix was generated, comparing the performance of models trained in one environment and tested in another (Figure 4). Higher predictive ability values (purple) indicate stronger predictive relationships, while lower values (yellow) suggest greater environmental variability or weaker genetic correlations.

**Figure 4.**
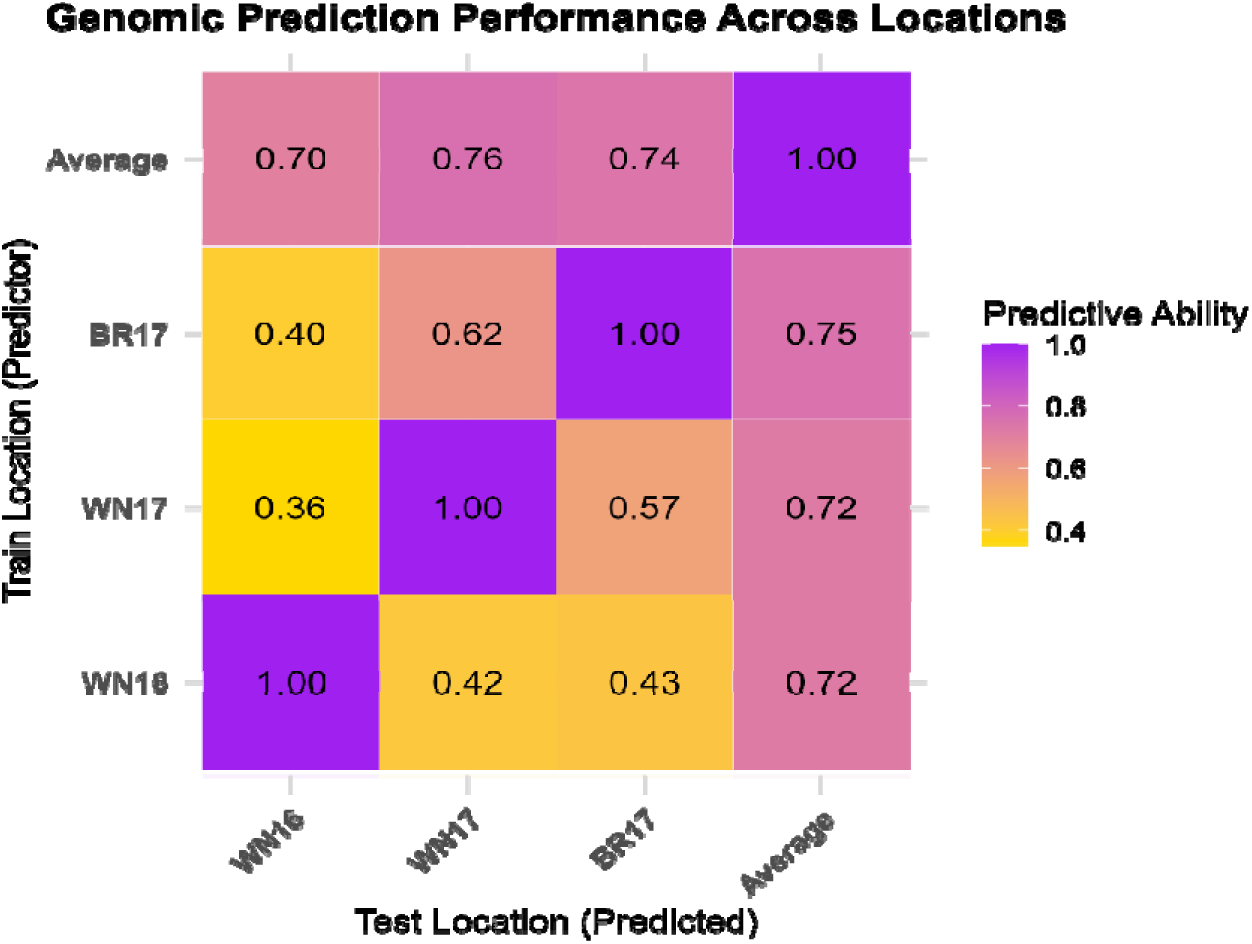
Predictive ability matrix for metribuzin tolerance across environments using GBLUP. Rows represent the training environment (predictor), and columns represent the validation environment (predicted). Values indicate Pearson correlation coefficients (r) between observed BLUEs and predicted genomic estimated GEBVs. Diagonal values represent within-environment prediction accuracy using cross-validation. Off-diagonal values represent cross-environment predictive ability. The “Average” column and row indicate the mean predictive ability across environments. Warmer colors (purple) indicate higher predictive ability, while lighter colors (yellow) indicate lower predictive ability.

Predictive ability (PA) varies across comparison. Using BR17 data as a training set and WN17 as a validation set gave a high prediction ability of 0.62, compared to 0.40 for WN16 as a validation set. WN16 showed consistently lower predictive ability across environments, with values of 0.42 and 0.43, indicating weaker agreement with the other locations. When the average values across all environments were used as the training set the prediction accuracy for all three environments was 0.70 to 0.76.

## 4 DISCUSSION

Metribuzin resistance is an important but understudied secondary trait for newly released wheat varieties in the SRWW region, due to its common use for weed control in wheat. A comprehensive understanding of the genetic mechanisms underlying metribuzin tolerance in our germplasm and environments will be beneficial for breeding resilient varieties capable of competing with weeds (Javid et al., 2017) by optimizing selection strategies for this trait in wheat breeding. The integration of phenotypic and genomic data remains a powerful tool for identifying key genetic factors influencing herbicide tolerance, and marker-based predictive efforts are likely to facilitate development of more tolerant varieties.

Previous studies demonstrated that metribuzin tolerance is governed by distinct genetic mechanisms across species. In soybean, tolerance is conferred by a single dominant gene; and in narrow-leafed lupin, tolerance arises from two semi-dominant loci with additive effects (Si et al., 2010). In field pea, a single QTL was identified, explaining 12–21% of the phenotypic variance, and appears to be associated with a non-target-site detoxification mechanism (Javid et al., 2017). A thorough understanding of genetic control and trait heritability is essential for optimizing selection strategies in metribuzin tolerance breeding. In this study we did not identify a major QTL for tolerance and most of the genotypes tested fell in the moderately resistant to resistant group. The broad-sense heritability estimates for metribuzin tolerance ranged from 0.54 to 0.76 across all site-years, indicating moderate genetic control. Prior studies have reported higher heritability values in more advanced breeding lines. Bhoite et al. (2019) documented broad-sense heritability estimates of 0.82, 0.95, and 0.92 for F5, F6, and F7 lines, respectively, reinforcing the notion that metribuzin tolerance is a moderately to highly heritable trait. The differences in heritability estimates may stem from the developmental stage of the lines assessed, as more advanced lines typically exhibit reduced environmental variance and greater genetic stability, leading to higher heritability values. Conversely, Villarroya et al. (2000) reported comparable broad-sense and narrow-sense heritability estimates of 0.52 and 0.23, respectively, in early-stage breeding lines, aligning with our findings. Their study attributed these estimates to the predominance of additive genetic variance, with the remaining variation influenced by dominance and epistatic interactions. This suggests that breeding methods using additive genetic variance can effectively develop superior genotypes.

Environmental factors influenced the observed variation in heritability estimates, particularly in WN16, where lower values were recorded. This decrease in heritability may be attributed to abnormal weather conditions including heavy rains and warm winter that impacted wheat stands and vernalization for that region during the trial period, which likely increased environmental noise and reduced the ability to capture genetic effects accurately. In this study, a fixed-effects GWAS model was implemented using the “GWAS()” function from the “rrBLUP” package, implemented as a least conservative model without principal components or kinship correction. Given the relatively high heritability of metribuzin tolerance, no major-effect SNPs surpassed the genome-wide significance threshold, which is consistent with polygenic architecture. This suggests that metribuzin tolerance is governed by multiple small-effect loci rather than a single, large-effect gene. The lack of significant associations is consistent with previous findings in complex quantitative traits influenced by environmental interactions. These results align with the moderate heritability estimates observed, further supporting the notion that metribuzin tolerance is a complex, polygenic trait.

The variability in genotypic responses across environments shows some evidence for GxE in metribuzin tolerance. However, the identification of consistently resistant genotypes such as AR04006-1 and AR05055-12-2 and recurrent susceptibility of lines like SCLA1084A1 and TX12D4896 suggests that screening methods can characterize lines with consistent levels of resistance. High correlation (ρ = 0.61) between BR17 and WN17, and even moderate correlations with WN16 (ρ = 0.41 – 0.43), further support the ability of methods to identify resistant genotypes. At the same time, these findings suggest that phenotypic selection based on data from a single environment, particularly one like WN16, may lead to inaccurate conclusions about true genetic tolerance. This challenge is common in breeding for abiotic stress tolerance, where environmental variability not only impacts trait expression but also reduces selection accuracy (Collard et al., 2005; Priess et al., 2020). Thus, multi-environment evaluation remains essential to evaluate genotypic tolerance.

Given the high overall level of resistance in southern SRWW germplasm, the ranking results from this study primarily serve to identify susceptible genotypes that should be excluded from breeding pipelines, rather than selecting new resistant parents for improvement. Marker-based selection methods should therefore seek to maintain this overall level of tolerance by minimizing the advancement of metribuzin-susceptible material. The combination of a lack of identified major QTL and high narrow-sense heritability suggest that genomic prediction is likely to be the best method for accomplishing this. Predictive ability (PA) varied considerably across environments, reflecting differences in site-specific environmental conditions. When BR17 was used as a training population for WN17, a strong across-environment PA of 0.62 was obtained, further supporting a shared genetic basis between these sites. In contrast, WN16 exhibited markedly lower predictive ability across environments (0.42–0.43). These observations emphasize that training populations encompassing multiple site–years provide more reliable across-environment predictions than models based on data from a single environment.

Polygenic traits, such as metribuzin tolerance, have a complex genetic architecture controlled by many small loci effect and strongly influenced by environmental variation. Due to the large number of allele effects, accurately predicting these traits with genomic prediction requires larger training populations covering many environments (Shahi et al., 2022). While results support making genomic selection a promising tool for breeding programs, recurring screening is required to further develop a continuing training population to predict breeding program lines.

## 5 CONCLUSION

While metribuzin tolerance was found to be moderately heritable, no major-effect loci were detected, suggesting that minor-effect loci collectively influence the trait. Genotypic responses across environments (WN16, WN17, BR17, and the combined dataset) revealed broad phenotypic variation. A consistent separation between highly tolerant and highly susceptible lines emphasizes not only the diversity in metribuzin response, but also the presence of lines with stable performance across variable environmental conditions, despite significant genotype-by-environment interactions.

The variation in predictive ability across environments evidences the strong influence of GxE interactions, which can change genetic signals and reduce model reliability. Despite these challenges, genomic selection demonstrated moderate accuracy, indicating its potential for improving metribuzin tolerance in wheat breeding. Integrating multi-environment testing with GS could enhance selection efficiency, particularly by targeting genotypes that are not only tolerant but also phenotypically stable across varying conditions. Incorporating these genotypes into crossing programs could enhance the development of metribuzin-tolerant cultivars with improved consistency across diverse growing conditions. Since high susceptibility to metribuzin occurs at a low frequency, genomic selection might be used at early stages to eliminate highly susceptible genotypes, rather than at a later stage to measure tolerance.

Given the widespread use of metribuzin in wheat production across major growing regions of the United States, including Kansas, Oklahoma, Texas, Colorado, Nebraska, and Louisiana, improving crop tolerance remains an important breeding goal. The development of metribuzin-tolerant cultivars can fortify the way to more reliable weed management options, reduce the risk of herbicide damage supporting stable yields across diverse environments.

## ACKNOWLEDGMENTS

The authors thank the reviewers and Associate Editor for their constructive comments and suggestions. This research was supported by the Louisiana Soybean and Grain Promotion Board.

## SUPPLEMENTAL MATERIAL

Supplementary Figure S1 presents the Spearman rank correlation matrix among site-years (BR17, WN16, and WN17) based on genotype BLUEs for metribuzin foliar damage. The figure provides additional information on the consistency of genotype rankings across environments and supports the assessment of environmental repeatability.

## Abbreviations

BLUE: Best Linear Unbiased Estimate
BLUP: Best Linear Unbiased Prediction
GBLUP: Genomic Best Linear Unbiased Prediction
GAWN: Gulf Atlantic Wheat Nursery
GWAS: Genome-Wide Association Study
GxE: genotype-by-environment interaction
H^2^: broad-sense heritability
h^2^: narrow-sense heritability
PA: predictive ability
SNP: single nucleotide polymorphism
SRWW: soft red winter wheat
GP: genomic prediction
QTL: quantitative trait loci
GEBV: genomic estimated breeding value

## REFERENCES

1. Albaugh, LLC. 2020. Metribuzin 4L herbicide label (EPA Reg. No. 42750-361). U.S. Environmental Protection Agency.

2. Baker, T.K., and Peeper, T.F. 1990. Differential tolerance of winter wheat (Triticum aestivum) to cyanazine and triazinone herbicides. Weed Technol. 4: 569–575.

3. Bhullar, M.S., Kaur, N., Kaur, P., and Gill, G. 2017. Herbicide resistance in weeds and its management. Agric. Res. J. 54: 436–444. doi:10.5958/2395-146X.2017.00085.0.

4. Bhoite, R., Si, P., Liu, H., Xu, L., Siddique, K.H.M., and Yan, G. 2019. Inheritance of pre-emergent metribuzin tolerance and putative gene discovery through high-throughput SNP array in wheat (Triticum aestivum L.). BMC Plant Biol. 19: 457. doi:10.1186/s12870-019-2070-x.

5. Collard, B.C.Y., Jahufer, M.Z.Z., Brouwer, J.B., and Pang, E.C.K. 2005. An introduction to markers, quantitative trait loci (QTL) mapping and marker-assisted selection for crop improvement: The basic concepts. Euphytica 142: 169–196. doi:10.1007/s10681-005-1681-5.

6. Cullis, B.R., Smith, A.B., and Coombes, N.E. 2006. On the design of early generation variety trials with correlated data. J. Agric. Biol. Environ. Stat. 11: 381–393. doi:10.1198/108571106X154443.

7. Endelman, J.B. 2011. Ridge regression and other kernels for genomic selection with R package rrBLUP. Plant Genome 4: 250–255. doi:10.3835/plantgenome2011.08.0024.

8. FAOSTAT. 2014. FAOSTAT Statistical Database: Wheat. Food and Agriculture Organization of the United Nations, Rome, Italy. Available at: https://www.fao.org/faostat/en.

9. Fahad, S., Hussain, S., Chauhan, B.S., Saud, S., Wu, C., Hassan, S., Tanveer, M., Jan, A., and Huang, J. 2015. Weed growth and crop yield loss in wheat as influenced by row spacing and weed emergence times. Crop Prot. 71: 101–108. doi:10.1016/j.cropro.2015.02.005.

10. Garcia-Baudin, J.M., Villarroya, M., Chueca, M.C., and Tadeo, J.L. 1990. Different tolerance of two cultivars of Triticum turgidum L. to metribuzin. Chemosphere 21: 223–230. doi:10.1016/0045-6535(90)90394-9.

11. Graymore, M., Stagnitti, F., and Allinson, G. 2001. Impacts of atrazine in aquatic ecosystems. Environ. Int. 26: 483–495. doi:10.1016/S0160-4120(01)00031-9.

12. Javid, M., Noy, D., Sudheesh, S., Forster, J.W., and Kaur, S. 2017. Identification of QTLs associated with metribuzin tolerance in field pea (Pisum sativum L.). Euphytica 213: 91. doi:10.1007/s10681-017-1878-4.

13. Kurya, B., Mia, M.S., Liu, H., and Yan, G. 2022. Genomic regions, molecular markers, and flanking genes of metribuzin tolerance in wheat (Triticum aestivum L.). Front. Plant Sci. 13: 842191. doi:10.3389/fpls.2022.842191.

14. Pilcher, W., Zandkamiri, H., Arceneaux, K., Harrison, S., and Baisakh, N. 2017. Genome-wide microarray analysis leads to identification of genes in response to herbicide metribuzin in wheat leaves. PLoS One 12: e0189639. doi:10.1371/journal.pone.0189639.

15. Priess, G.L., Norsworthy, J.K., Roberts, T.L., Spurlock, T.N., and Gbur, E.E. 2020. Soybean growth and incidence of soil-borne fungi as influenced by metribuzin. Agron. J. 112: 5132–5142. doi:10.1002/agj2.20396.

16. Runyan, T.J., McNeil, W.K., and Peeper, T.F. 1982. Differential tolerance of wheat (Triticum aestivum L.) cultivars to metribuzin. Weed Sci. 30: 94–97. doi:10.1017/S0043174500026242.

17. Shahi, D., Guo, J., Pradhan, S., Khan, J., Avci, M., Khan, N., McBreen, J., Bai, G., Reynolds, M., Foulkes, J., and Babar, M.A. 2022. Multi-trait genomic prediction using in-season physiological parameters increases prediction accuracy of complex traits in US wheat. BMC Genomics 23: 298. doi:10.1186/s12864-022-08487-8.

18. Shaner, D.L. (ed.). 2014. Herbicide Handbook. 10th ed. Weed Science Society of America, Lawrence, KS.

19. Si, P., Pan, G., and Sweetingham, M. 2010. Semi-dominant genes confer additive tolerance to metribuzin in narrow-leafed lupin (Lupinus angustifolius L.) mutants. Euphytica 177: 411–418. doi:10.1007/s10681-010-0278-9.

20. Siddiqui, I., Bajwa, R., Huma, Z., and Javaid, A. 2010. Effect of six problematic weeds on growth and yield of wheat. Pak. J. Bot. 42: 2461–2471.

21. Singh, R., Hager, A., Lancaster, S., Norsworthy, J.K., Gage, K., Johnson, W., Young, B., Stephenson, D., Bond, J., Bradley, K., Jhala, A., Essman, A., Steckel, L.E., Mueller, T.C., Sprague, C., Legleiter, T., Werle, R., Ikley, J., Jha, P., and Jugulam, M. 2025. Optimizing metribuzin rates for herbicide-resistant Amaranthus weed control in soybean. Weed Technol. 39: e97. doi:10.1017/wet.2025.10047.

22. Trebst, A. 2007. The mode of action of triazine herbicides in plants. Photosynth. Res. 93: 223–229. doi:10.1007/s11120-007-9178-y.

23. Villarroya, M., Escorial, M.C., Garcia-Baudin, J.M., and Chueca, M.C. 2000. Inheritance of tolerance to metribuzin in durum wheat. Weed Res. 40: 293–300. doi:10.1046/j.1365-3180.2000.00188.x.

24. Yu, J., Pressoir, G., Briggs, W.H., Bi, I.V., Yamasaki, M., Doebley, J.F., McMullen, M.D., Gaut, B.S., Nielsen, D.M., Holland, J.B., Kresovich, S., and Buckler, E.S. 2006. A unified mixed-model method for association mapping that accounts for multiple levels of relatedness. Nat. Genet. 38: 203–208. doi:10.1038/ng1702.

