## Supplemental Figure 1 for "Genome-Wide Markers Predict Metribuzin Tolerance in Southern Soft Red Winter Wheat"

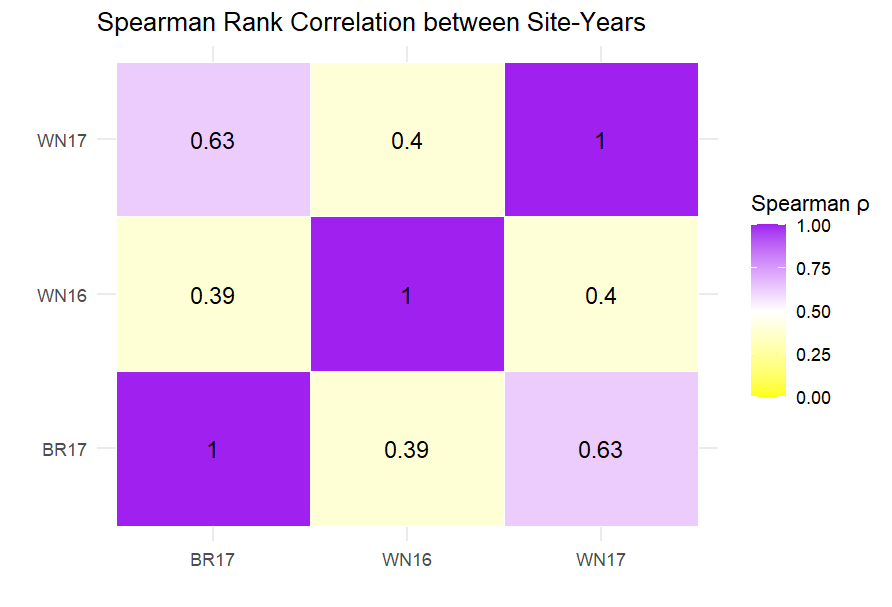

**Supplementary Figure 1.** Spearman rank correlation matrix for metribuzin tolerance across three site-years (BR17, WN16, and WN17). Values represent Spearman correlation coefficients (ρ) calculated from genotype BLUEs for foliar damage scores. Higher correlations indicate greater consistency in genotype rankings across environments. The strongest correlation was observed between BR17 and WN17 (ρ = 0.63), whereas lower correlations were observed between BR17 and WN16 (ρ = 0.39) and between WN16 and WN17 (ρ = 0.40). Darker purple colors indicate stronger positive correlations, while lighter yellow colors indicate weaker correlations.
